# Interaction of RSC chromatin remodelling complex with nucleosomes is modulated by H3 K14 acetylation and H2B SUMOylation *in vivo*

**DOI:** 10.1101/2020.03.02.972562

**Authors:** Neha Jain, Davide Tamborrini, Brian Evans, Shereen Chaudhry, Bryan J. Wilkins, Heinz Neumann

**Affiliations:** Department of Structural Biochemistry, Max-Planck-Institute of Molecular Physiology, Otto-Hahn-Strasse 11, 44227 Dortmund, Germany; Department of Chemistry and Biochemistry, Manhattan College, 4513 Manhattan College Parkway, Bronx, NY 10471; Department of Chemical Engineering and Biotechnology, University of Applied Sciences Darmstadt, Stephanstrasse 7, 64295 Darmstadt

**Keywords:** Genetic Code Expansion, Unnatural Amino Acids, Photo-crosslinking, Chromatin Remodelling, SUMOylation, Lysine acetylation

## Abstract

Chromatin remodelling complexes are multi-subunit nucleosome translocases that reorganize chromatin in the context of DNA replication, repair and transcription. A key question is how these complexes find their target sites on chromatin. Here, we use genetically encoded photo-crosslinker amino acids to map the footprint of Sth1, the catalytic subunit of the RSC (remodels the structure of chromatin) complex, on the nucleosome in living yeast. We find that the interaction of the Sth1 bromodomain with the H3 tail depends on K14 acetylation by Gcn5. This modification does not recruit RSC to chromatin but mediates its interaction with neighbouring nucleosomes. We observe a preference of RSC for H2B SUMOylated nucleosomes *in vivo* and show that this modification moderately enhances RSC binding to nucleosomes *in vitro*. Furthermore, RSC is not ejected from chromatin in mitosis, but its mode of nucleosome binding differs between interphase and mitosis. In sum, our *in vivo* analyses show that RSC recruitment to specific chromatin targets involves multiple histone modifications most likely in combination with other components such as histone variants and transcription factors.

**Key Points:**

- *In vivo* photo-crosslinking reveals the footprint of the ATPase subunit of RSC on the nucleosome.
- RSC binds to H3 K14ac nucleosomes via the C-terminal bromodomain of its ATPase-subunit Sth1.
- RSC preferentially localizes to H2B-SUMOylated nucleosomes.

## Introduction

Storage and accessibility of genetic information are two conflicting requirements that a cell must balance. While the DNA must be compacted to meet the space limitations of the nucleus, access to its information content for transcription, repair, and replication processes must be ensured. Eukaryotes accomplish this by packaging their DNA in chromatin, however, densely packed chromatin territories restrict the accessibility of the underlying DNA [1]. Hence, to facilitate access to DNA, chromatin has evolved highly malleable properties to meet the demands for dynamic changes [2–5].

Chromatin remodelling enzymes use ATP hydrolysis to rearrange nucleosomes to enable other factors to access DNA [6]. Posttranslational modifications (PTMs) of histones and histone variants either modulate the stability and DNA-binding properties of the nucleosome or signal the recruitment of machinery that initiates the transcription, replication or repair of DNA [5]. The turnover of most histone PTMs is rapid, making these processes very dynamic.

Common to all families of chromatin remodelers are an affinity for nucleosomes, the ability to recognize histone PTMs via specialized domains, and a DNA-dependent ATPase domain that translocates DNA relative to the histone octamer. Apart from that, remodelers differ significantly in subunit composition, specificity for histone modifications, the processes in which they are involved, and whether they promote chromatin opening or closing. How these enzymes work, how their activity is regulated, and how they are recruited to specific loci is currently being actively investigated.

The RSC complex is an abundant, essential chromatin remodelling complex of the SWI/SNF family in budding yeast [7]. RSC is involved in transcription [8–12], chromosome segregation [13], replication [14], and the response to DNA damage [15, 16].

Recently, the high-resolution structure of the RSC complex bound to a nucleosome has been solved by cryo-electron microscopy, revealing three flexibly connected parts [17–19]. The motor domain of the Sth1 subunit binds at superhelical location (SHL) +2, from where it translocates the DNA one base pair at a time into the direction of the dyad, possibly creating a loop that propagates around the nucleosome [20]. The ARP module, which couples DNA translocation and ATPase activity [21], connects the motor domain to the substrate recognition module (SRM). The latter contains DNA-binding Zn-cluster domains, five bromodomains, a histone-tail binding BAH domain and the nucleosome-binding C-terminal tail of Sfh1 [17–19]. Due to the flexible tethering of these domains, their structure, substrate preference, and interaction with the nucleosome remained largely unresolved. The only established lysine acetylation site on histones recognized by RSC is H3 K14ac by the bromodomains of Sth1 and Rsc4 [22–24].

Here, we use quantitative *in vivo* crosslinking with genetically encoded photo-activatable crosslinker amino acids to reveal the footprint of the Sth1 subunit of RSC on the nucleosome. The interaction of Sth1 with the N-terminal H3 tail depends on the presence of H3 K14, which is recognized by the C-terminal bromodomain of Sth1 upon acetylation [22]. We further show that Sth1 preferentially crosslinks to the SUMOylated form of H2B, suggesting that H2B SUMOylation acts in the context of RSC remodelling.

## Materials and Methods

### Chemicals

Anti-HA antibody was obtained from Abcam (ab9110), pBPA from Chem-Impex. Other materials and media were used as reported previously [25].

### Yeast strains and plasmids

Histone amber mutants were created by QuikChange mutagenesis on pRS426 plasmids encoding a C-terminally HA-tagged histone under the control of its native promoter and verified by sequencing the entire ORF [25]. pCDF-ScNap1 was created by cloning the *NAP1* ORF in pCDF-DUET using restriction sites BamHI/SalI. pQE80-His_14_-SUMO-H2B was created by cloning the *Xenopus* H2B-ORF lacking the first five codons in pQE80-His_14_-SUMO (a kind gift of Dirk Görlich, Göttingen) using restriction sites KpnI/SpeI. Histone octamers containing SUMOylated H2B were produced using a plasmid encoding for all four human core histones in which the ORF of H2B was replaced by the ORF of pQE80-His_14_-SUMO-H2B. Plasmids encoding multiple repeats of Widom-601 sequences were gifts of Fabrizio Martino and Daniela Rhodes. Yeast strains used in this study are listed in Table 1. BY4741 Sth1-3myc was obtained by integrating three myc-epitopes followed by the HIS3MX6 cassette of pYM5 [26] prior to the translational stop codon of the *STH1* gene using standard PCR-based tagging. BJ3505-Rsc2-TAP was created in analogous fashion by integrating CBP-TEV-protA at the end of the RSC2 ORF using plasmid pBS1479 [27]. Successful integration was verified by PCR and Western blot. BY4741 Δ*siz1* Δ*siz2* was created in two sequential rounds of genomic integrations. First, the *kanMX* box was used to replace the *SIZ1* locus in BY4741 by a PCR product amplified from pUG6 [28] with flanking sequences homologous to the *SIZ1* 5’- and 3’-UTR. Successful replacement was verified by PCR of geneticin resistant clones. Next, *SIZ2* was replaced by genomic integration of the *hphNT1* box amplified from pYM24 [29] in the same manner using hygromycin B selection.

**Table 1.**
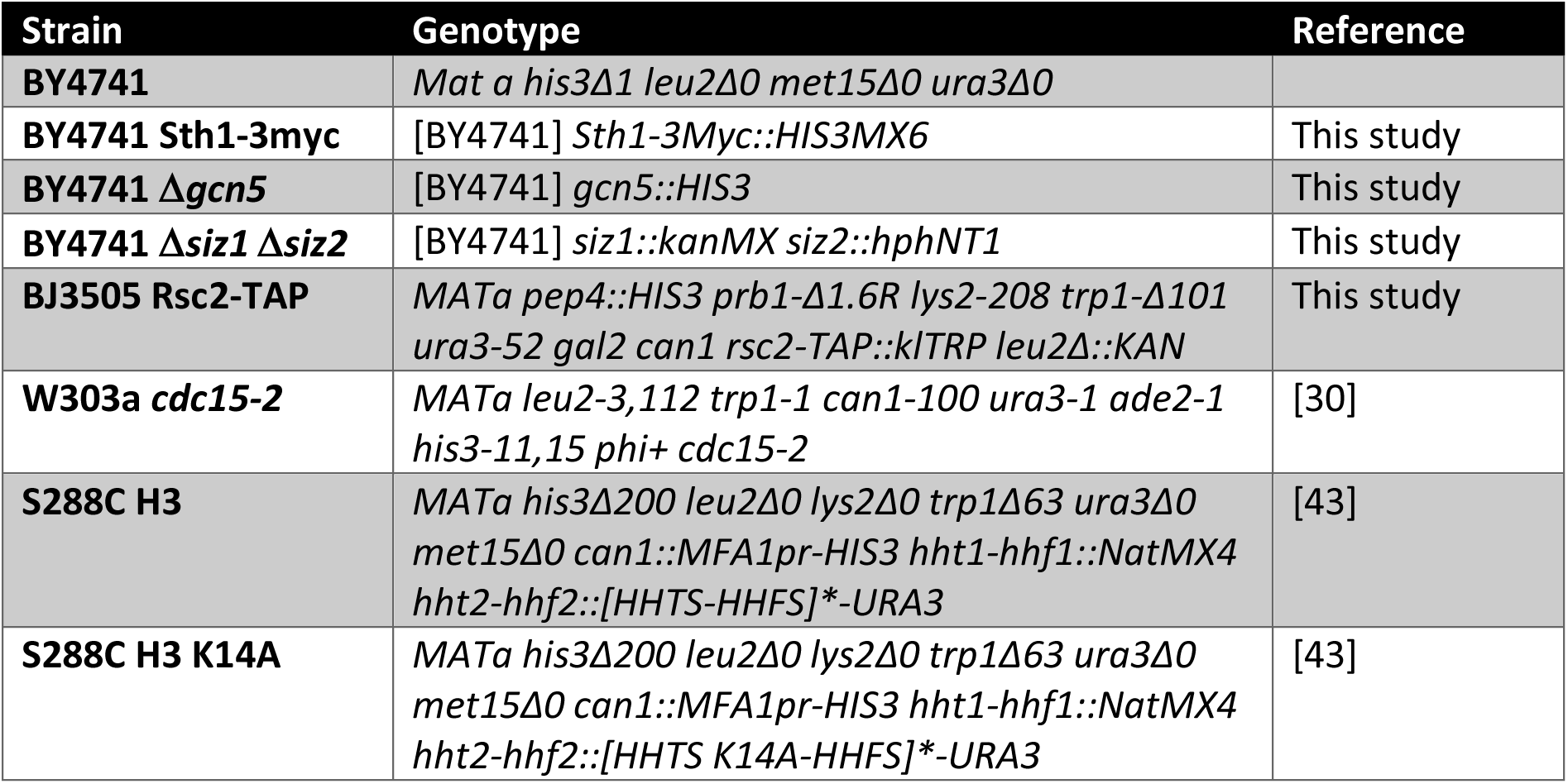
Yeast strains.

BY4741 Δ*gcn5* was established through genomic integration of the *HIS3* gene at the *GCN5* locus. Standard PCR amplification of the pRS303 plasmid was performed to amplify the *HIS3* region with flanking sequences homologous to the *GCN5* 5’- and 3’-UTR. Genomic integration was selected with SC-His dropout agar and surviving colonies were verified by PCR.

### Photo-crosslinking of living yeast

Yeast photo-crosslinking experiments were carried out as described [25]. Briefly, yeasts were transformed with plasmids to encode pBPA-containing histones and grown in SC-Ura/Leu with 1 mM pBPA. For crosslinking, 12 OD units were harvested, irradiated (365 nm) for 15 min on ice, proteins extracted by alkaline lysis and precipitated with TCA. Crosslinks were analysed by 3-8% Tris-acetate SDS-PAGE and Western blot.

Blots were developed by measuring chemiluminescence (ECL Select, GE Healthcare) with an imaging system (Celvin S, Biostep). Quantification of band intensities were performed with Fiji software on raw inverted tiff images of Western blots. Therefore, the image was rotated so that lanes ran horizontally. A box was drawn across an individual lane using the “rectangle” tool and grey density analysed by “Plot profile”. The peak corresponding to the protein band was traced using the “free hand selection” tool and the peak area analysed using “measure”.

### Yeast cell cycle synchronization

Cell cycle synchronizations were performed in W303a cells with a temperature-sensitive *cdc15-2* allele [30] containing plasmids encoding H2B or H3 pBPA-mutants [31]. An overnight culture of 4 ml in SC-Ura/Leu minimal medium (supplemented with 1 mM pBPA) was diluted to an OD600 = 0.2 in 50 ml YPD and incubated for 2 h, with shaking at 200 rpm, at 25°C. Upon reaching OD600 = 0.5, cells were shifted for 2 h to 37°C with shaking to arrest cells in telophase [32]. Following complete arrest (as determined by FACS analysis [33]), cells were released into 50 ml YPD at an OD600 = 0.5 at 25°C. Samples were taken at indicated time points, crosslinked and processed as described above.

### RSC purification

RSC complex was purified by tagging the C-terminus of the Rsc2 subunit with a TAP tag in BJ3505. RSC was purified essentially as described [27].

### Nucleosome Reconstitution

#### Purification of Octamers

DNA-Sequence from yeast *SMT3* was fused in frame to the K6 of *Xenopus* H2B and cloned via Gibson Assembly into a pDUET plasmid for polycistronic coexpression (a gift from A. Musacchio) [34]. The final construct containing His_6_H3.1-H4-His_6_H2A1B-SUMO-H2B, was used to transform Rosetta2^®^ *E. coli* cells. Expression and purification were carried out essentially as described [35]. After elution from the Ni-NTA column, the octamer-containing fractions were concentrated with an Amicon 10k. The His-tag was removed with 0.2 mg/ml GST-PreScission protease for 12 hours at 4°C. For the purification of deSUMOylated octamers, Ubl1 protease was also added to the concentrated octamers to a final concentration of 0.2 mg/mL. The digested octamers were further purified by gel-filtration (Superdex 200 10/300 GL, GE Healthcare) in 10 mM HEPES pH 7.4, 2.0 M NaCl, 0.5 mM EDTA, 1 mM TCEP. Fractions containing the octamers were pooled and used for nucleosome reconstitution.

#### DNA purification

DNA for nucleosome reconstitution was obtained from a pUC18 plasmid containing 25 repeats of Widom 601 DNA with 25 bp overhang on both sides (a gift from D. Rhodes). The 197 bp array was digested out from the plasmid with AvaI and concentrated to 1 mg/mL. The plasmid backbone was precipitated by centrifugation at 14000 g for 30 minutes after 12 hours incubation at 4°C in 10 mM Tris pH 8, 700 mM NaCl, 10 mM MgCl_2_, 1 mM EDTA, 6.5% PEG 8000. The DNA in the supernatant was precipitated with isopropanol and used for nucleosome reconstitution. For the nucleosomes employed in the BLI experiments, we used the 167 bp 601 Widom DNA, which was purified with a similar protocol.

#### Nucleosome Reconstitution

Increasing concentration of octamers were titrated against a fixed amount of 197 bp Widom DNA to accurately determine the best ratio of protein and DNA. To reconstitute the final nucleosomes, a dialysis bag containing 1 mL solution of 3.0 µM DNA and 3.0 µM octamers was inserted into a second dialysis bag containing 50 mL of 10 mM HEPES pH 7.4, 2.0 M NaCl, 0.5 mM EDTA, 1 mM TCEP, and the resulting double bag was dialysed at 4°C for 20 h against 5 L 10 mM HEPES pH 7.4, 35 mM KOAc, 0.5 mM EDTA, 5 mM 2-BME. The quality of the reconstitutions was assessed by 6% Native PAGE (Acrylamide/Bis-acrylamide 49:1 ratio) stained with Gel Red^®^.

### Purification of scNap1

His-tagged scNap1 was expressed and purified in Rosetta2^®^ *E. coli* cells essentially as described [36]. After Ni-NTA elution, Nap1 containing fractions were concentrated to 2 mg/mL and loaded on a Superdex 200 10/300 GL gel-filtration column (GE Healthcare) pre-equilibrated with 25 mM HEPES pH 7.4, 10% glycerol, 300 mM KCl, 2 mM MgCl_2_, 1 mM DTT. 6xHis-scNap1 containing fractions were pooled, concentrated to 24 µM and stored at –80°C.

### Nucleosome Remodelling Assay

All remodelling reactions were performed at 30°C in 15 µl volumes containing 10 nM RSC complex, 60 nM NCPs, and 1.5 µM scNap1 in reaction buffer (10 mM HEPES pH 7.4, 250 mM KOAc, 2 mM MgCl_2_, 0.6 mM ATP, 100 µg/mL BSA). Reactions were quenched with 600 ng lambda DNA and glycerol (5% final concentration) after 1, 3, 7, 15, or 30 minutes and kept at 30°C for additional 15 min after quenching before being shifted to 4°C. To address the high velocity of the reaction, the time 0 reactions were incubated for 1 minute with 0.6 mM ADP instead of ATP and quenched as previously described.

To assess NCP remodelling, 6 µl of remodelling reactions were loaded onto 6% Native PAGE gel (Acrylamide/Bis-acrylamide 49:1 ratio) in 0.4x TBE, run in 0.4x TBE buffer at 4°C for 90 min at 150 V, and stained with Gel Red^®^. Band intensities were analysed and quantified with ImageJ.

### RSC-NCP binding kinetic

After purification, the RSC complex was incubated with 10-fold molar excess of N-hydroxysuccinimidobiotin at room temperature and after 90 min the reaction was quenched with 25 mM Tris pH 8.0 for 10 minutes before loading the biotinylated RSC complex onto a MiniQ column pre-equilibrated in RSC buffer 200 (5% glycerol, 10 mM HEPES pH 8.0, 200 mM KOAc, 0.5 mM EDTA, 0.5 mM TCEP) and eluted with linear gradient of increasing concentration of RSC buffer 1 M KOAc. The RSC complex typically eluted at 700 mM KOAc. The presence of all subunits was confirmed by SDS-PAGE.

The RSC complex was loaded onto Streptavidin (SA) biosensors which were quenched with free biotin after loading. Binding and dissociation buffers only differ in the presence/absence of free NCPs and both contained 5% glycerol, 10 mM HEPES pH 8.0, 80 mM KOAc, 0.5 mM EDTA, 2.5 mM MgCl_2_, 0.5 mM AMP-PNP, and 0.3 mg/mL BSA.

### RSC ATPase Activity

To determine the Michaelis-Menten constant of RSC complex on both WT and SUMOylated NCPs reconstituted with the 197 bp 601 Widom DNA, we evaluated the ATPase activity of Sth1 with an ADP-Glo™Assay (Promega). The reactions were performed at 30°C in 30 µl containing 5% glycerol, 10 mM HEPES pH 7.4, 120 mM KOAc, 150 µg/mL BSA, 2.5 mM MgCl_2,_ 0.5 mM ultrapure ATP, 0.5 mM TCEP, and 10 nM RSC. After 5, 10, and 15 minutes 5 µl were quenched with ADP-Glo™ Reagent and analysed as recommended by the supplier. All measurements were performed in triplicate and analysed with GraphPad.

## Results

### In vivo cross-linking survey of the nucleosome

Genetic code expansion allows for the incorporation of unnatural amino acids in response to amber (UAG) stop codons in a variety of cells and organisms [37]. We have established the incorporation of photo-activatable crosslinker amino acids in histones and chromatin-interacting proteins to study the dynamics of chromatin in living yeast [31, 38]. The crosslinking reaction follows a long-wavelength ultraviolet light (365 nm) inducible, radical mechanism that results in the formation of binary covalent adducts that can be quantitated by Western blot [39].

To map the interactome of the nucleosome in living yeast, we created a library of more than one hundred amber mutants covering the surface-exposed residues of the nucleosome. We incorporated *p*-benzoyl-L-phenylalanine (pBPA) in response to amber stop codons by genetic code expansion for each individual mutant and analysed crosslink products formed upon irradiation by SDS-PAGE and Western blot against the HA-epitope on the histone (Fig. S1 & 2). Crosslink scans of the nucleosomal surface revealed differential binding patterns across each of the histones, highlighting the viability of interactome mapping in the living nucleus. We then asked if this approach could be employed to characterize individual chromatin binding proteins, specifically nucleosome bound chromatin remodelling complexes.

### Footprint of Sth1, the catalytic subunit of RSC, on the nucleosome

In order to probe the footprint of the catalytic subunit of RSC, Sth1, on the nucleosome surface we selected 58 histone pBPA mutants from the crosslinking survey that had produced a band of appropriate combined mass for a crosslink product of a histone with Sth1 (approximately 170 kDa). To test whether these bands indeed result from crosslinking to Sth1 we performed the same crosslinking reaction in strains with or without a C-terminal 3xmyc-tag on Sth1 (Fig. 1, Fig. S3-6). The epitope tag leads to a slower migration of the corresponding band of the crosslink product in Western blots. This analysis identified nine positions on the nucleosome core and H3 tail that interact with Sth1 (Fig. 1A). We detected two further positions in the H3 tail (T6 and T11), when we precipitated the crosslink products with anti-myc antibody beads (against myc-tagged Sth1) prior Western blot analysis with anti-HA antibodies (Fig. 1B). This confirmed the ability of pBPA at these positions to crosslink to Sth1. We mapped these positions on the structure of the nucleosome core particle to visualize the footprint of Sth1 (Fig. 1C).

**Figure 1.**
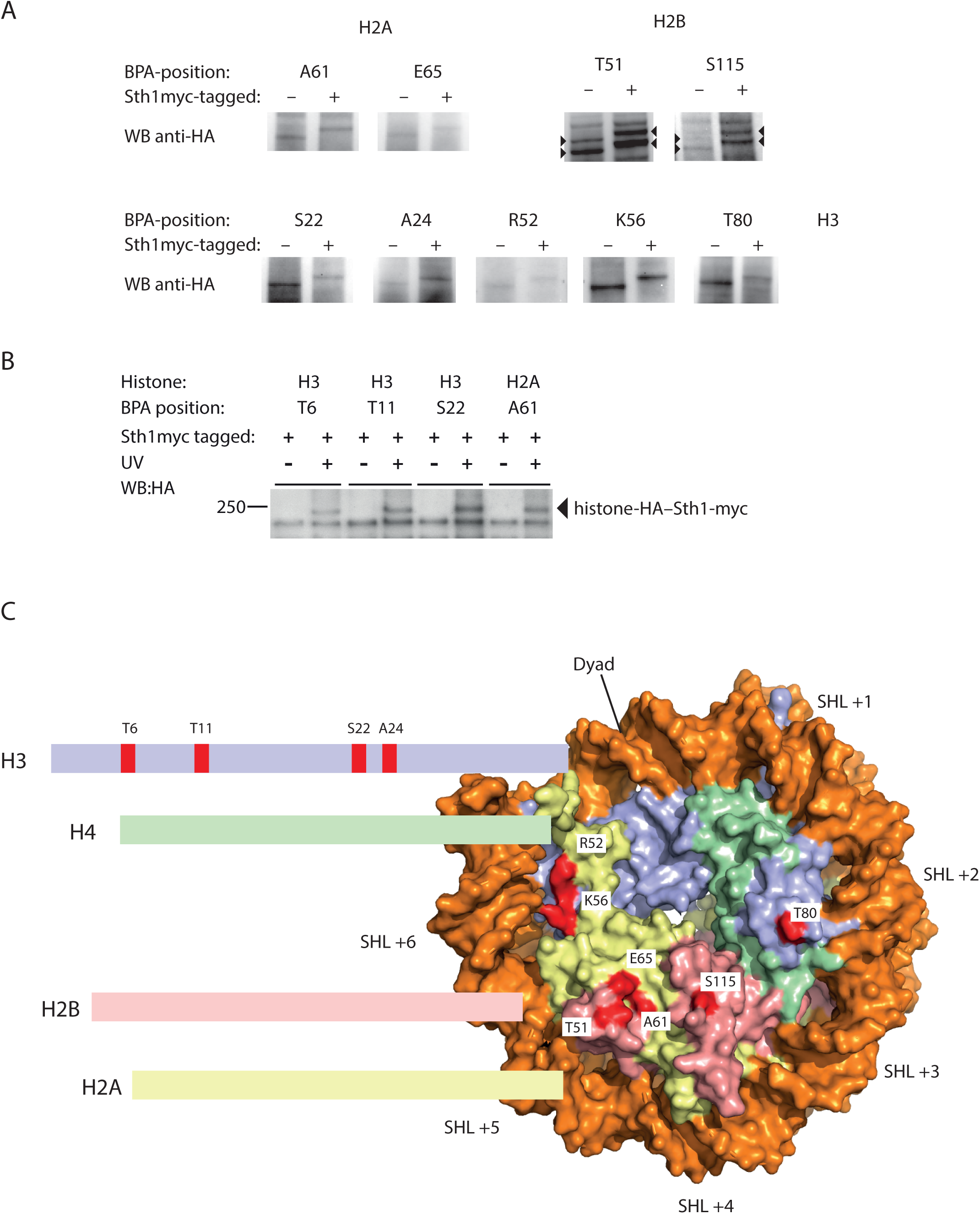
Mapping the interaction surface of Sth1 on the nucleosome *in vivo*. A) Yeast cells (wild-type or Sth1-3myc) expressing histones with pBPA at the indicated position were UV-irradiated and crosslink products analysed by SDS-PAGE and Western blot using anti-HA antibodies recognising the pBPA-containing histone. The shift in mobility resulting from the myc-tag identifies a crosslink product with Sth1. H2B-Sth1 crosslinks appear as a double band due H2B SUMOylation (arrow heads). B) Crosslink reactions from positions in the H3 tail performed in Sth1-3myc cells were subjected to immunoprecipitation with anti-myc antibody beads prior to analysis by Western blot with anti-HA antibodies. C) Graphical representation of the positions identified in A and B on the structure of the nucleosome. Figure was prepared using pdb-file 1ID3 and PyMol v1.7.6.6.

### Binding of the H3 tail by Sth1 depends on H3 K14

We hypothesised that the C-terminal bromodomain of Sth1 might mediate the interaction with the H3 tail because H3 K14 acetylation enhances nucleosome binding by RSC [40] and a recent co-crystal structure revealed an extensive interface between the Sth1 bromodomain and an H3 tail peptide [22]. We therefore mutated H3 K14 to alanine on the same H3 copy containing pBPA. The mutation interfered with crosslinking of H3 T6pBPA and T11pBPA to Sth1, while crosslinking from H3 S22pBPA was only partially affected by K14A, suggesting that K14 is required for the interaction of Sth1 with the tip of the H3 tail (Fig. 2A).

In order to demonstrate that acetylation of H3 K14 is essential for this interaction, we deleted the gene encoding lysine acetyltransferase Gcn5, the enzyme responsible for the deposition of H3 K14ac [41, 42]. This indeed abolished crosslinking between the H3 tail and Sth1, much like the H3 K14A mutation (Fig. 2B).

**Figure 2.**
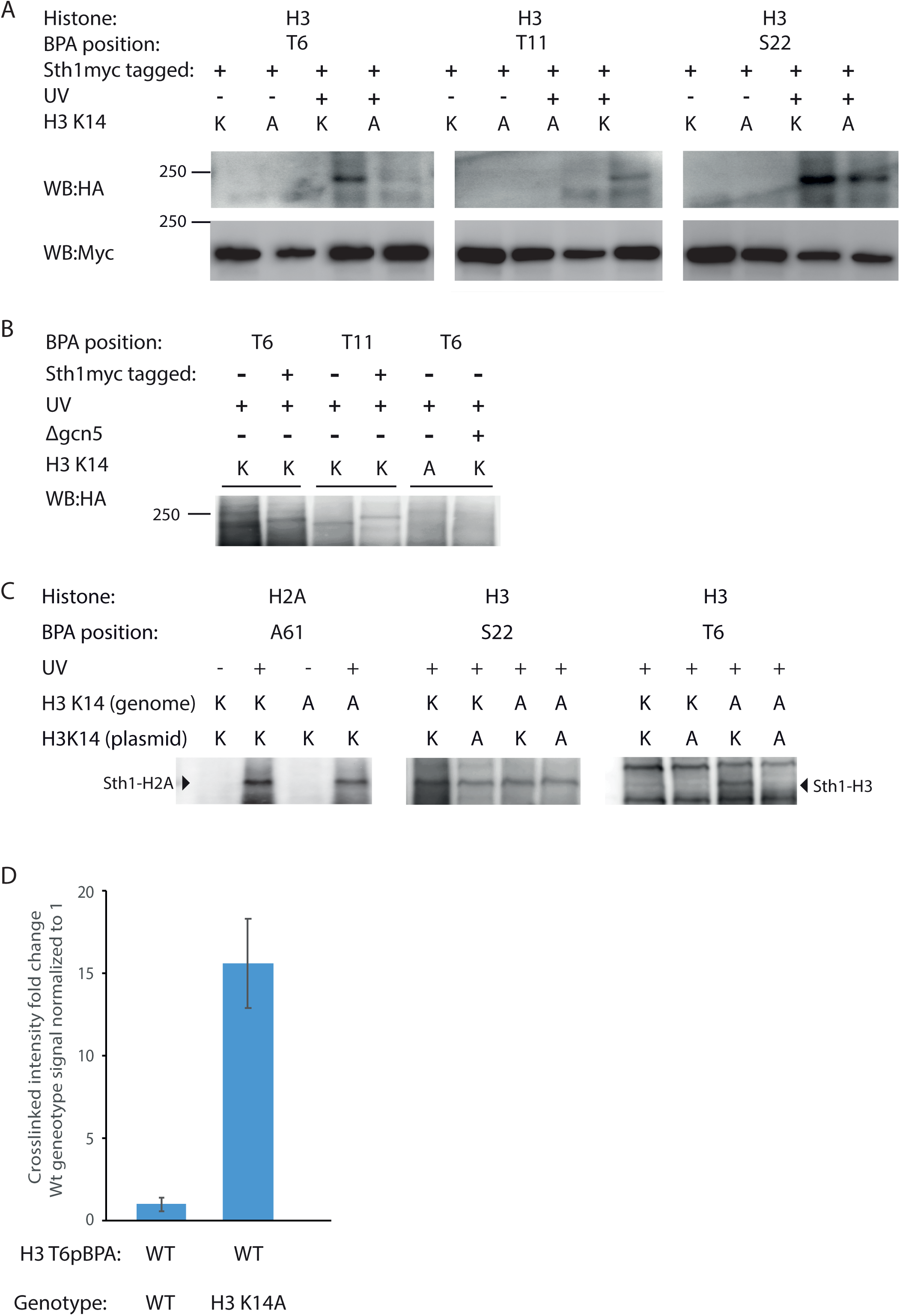
Crosslinking of the H3 tail to Sth1 is regulated by H3 K14ac. A) Crosslinking from positions in the H3 tail in Sth1-3myc cells. Anti-myc immunoprecipitates were analysed by Western blot with anti-HA antibodies. B) Deletion of *gcn5* interferes with H3 T6pBPA crosslinking to Sth1. WCEs of crosslinked samples were irradiated with UV-light and whole cell extracts analysed by Western blot using anti-HA antibodies. C) Effect of mutating endogenous H3 K14 to alanine on crosslinking to Sth1. Yeasts (wild-type or H3 K14A) expressing H3 T6pBPA, H3 S22pBPA or H2A A61pBPA with or without K14A mutation were analysed as in B. D) Quantitative comparison of crosslinking efficiencies from H3 T6pBPA in wild-type and H3 K14A yeasts. Error bars are standard deviations of five independent experiments.

Next, we asked whether H3 K14ac serves to recruit RSC to nucleosomes. Therefore, we performed crosslinking experiments from H2A A61pBPA–nucleosomes in yeast with or without a genomic H3 K14A mutation (Fig. 2C, left panel) [43]. We observed that crosslinking of Sth1 was only slightly reduced by the K14A mutation. Hence, recruitment of RSC to chromatin does not require H3 K14ac, otherwise the crosslinking efficiency from this position would have been reduced in the mutant background.

This is consistent with the micromolar concentration of nucleosomes in the yeast nucleus (30,000 nucleosomes in a volume of 3 fL) being approximately one thousand times greater than the dissociation constant (K_D_) of RSC–nucleosome complexes [44]. Therefore, increasing the affinity of RSC for nucleosomes by histone modifications is not expected to enhance the level of saturation (Θ) of RSC with nucleosomes because Θ is hardly affected by changes in ligand concentration if their concentration is greater than ten times K_D_.

However, H3 K14ac may control which nucleosomes are bound by RSC. In this case, mutation of H3 K14 in nucleosomes without the crosslinker (i.e. in the genomic copy of the H3 gene) should shift RSC binding to crosslinker-containing nucleosomes which still possess H3 K14ac.

Therefore, we performed crosslinking experiments from H3 S22 in the background of a yeast strain bearing the K14A mutation in the genomic copy of H3 (Fig. 2C, middle panel). If H3 K14ac recruits RSC to nucleosomes, this mutation should increase the crosslinking efficiency because the crosslinker-containing nucleosomes are the only ones with an intact H3 K14 residue. However, crosslinking from H3 S22pBPA was not affected by genomic H3 K14A mutation, hence, H3 K14ac does not control recruitment of RSC.

Finally, we asked whether the Sth1 bromodomain exclusively interacts with H3 tails that are part of the nucleosome bound by RSC or whether H3 tails from neighboring nucleosomes are also substrates. Therefore, we performed crosslinking experiments from H3 T6pBPA in the genetic background of H3 K14A cells (Fig. 2C, right panel). If the bromodomain of Sth1 only interacts with H3 tails that are part of the same nucleosome that the complex is bound to, the mutation of the endogenous H3 should not affect the crosslinking efficiency. However, if Sth1 interacts with H3 tails of neighbouring nucleosomes, the mutation would abrogate the competition with their histone tails and therefore increase crosslinking. Indeed, the K14A mutation of the endogenous H3 allele increased crosslinking between Sth1 and H3 T6pBPA fifteenfold (Fig. 2D), strongly indicating that the bromodomain of Sth1 is able to interact with acetylated H3 tails of other nucleosomes.

### RSC preferentially interacts with SUMOylated H2B in vivo

Crosslink products of Sth1 to histones H2A and H3 each migrated as a single band in Western blots with an apparent molecular mass of about 180 kDa (Fig. 1). H2B-Sth1 crosslink products (from positions T51 and S115), however, showed a second band shifted by approximately 10 kDa to a higher apparent mass (Fig. 1). Initially, we hypothesised that this mass shift is a result of H2B K123 ubiquitination, the major site of H2B ubiquitination in budding yeast [45]. However, the crosslinking pattern of ubiquitination-deficient mutant H2B T51pBPA K123R was indistinguishable from that of H2B T51pBPA (Fig. 3A). Hence, we tested additional H2B sites known to be ubiquitinated (K46, K49 and K111) [46], which also did not change the crosslinking pattern (Fig. 3A). Next, we analysed the impact of mutating combinations of K6, K7, K16 and K17 in H2B T51pBPA to Arg on crosslinking to Sth1 (Fig. 3B). These are the major sites of H2B SUMOylation in *S. cerevisiae* [47]. Indeed, in the absence of these lysine residues, we observed only a single H2B-Sth1 crosslink product. Interestingly, mutating either pair of lysine residues in the H2B N-terminus (K6/7 or K16/17) was sufficient to abolish the slower migrating band. Accordingly, mutations of the same sites have previously been shown to abolish H2B SUMOylation [47]. Finally, when we analysed a strain lacking E3 SUMO ligases Siz1 and Siz2 required for H2B SUMOylation [47], the crosslink reaction produced only a single H2B-Sth1 band (Fig. 3C), unambiguously confirming the upper band to be the SUMOylated form of H2B. Densitometric quantification of both bands indicates that 20-30% of H2B that interacts with Sth1 is SUMOylated.

**Figure 3.**
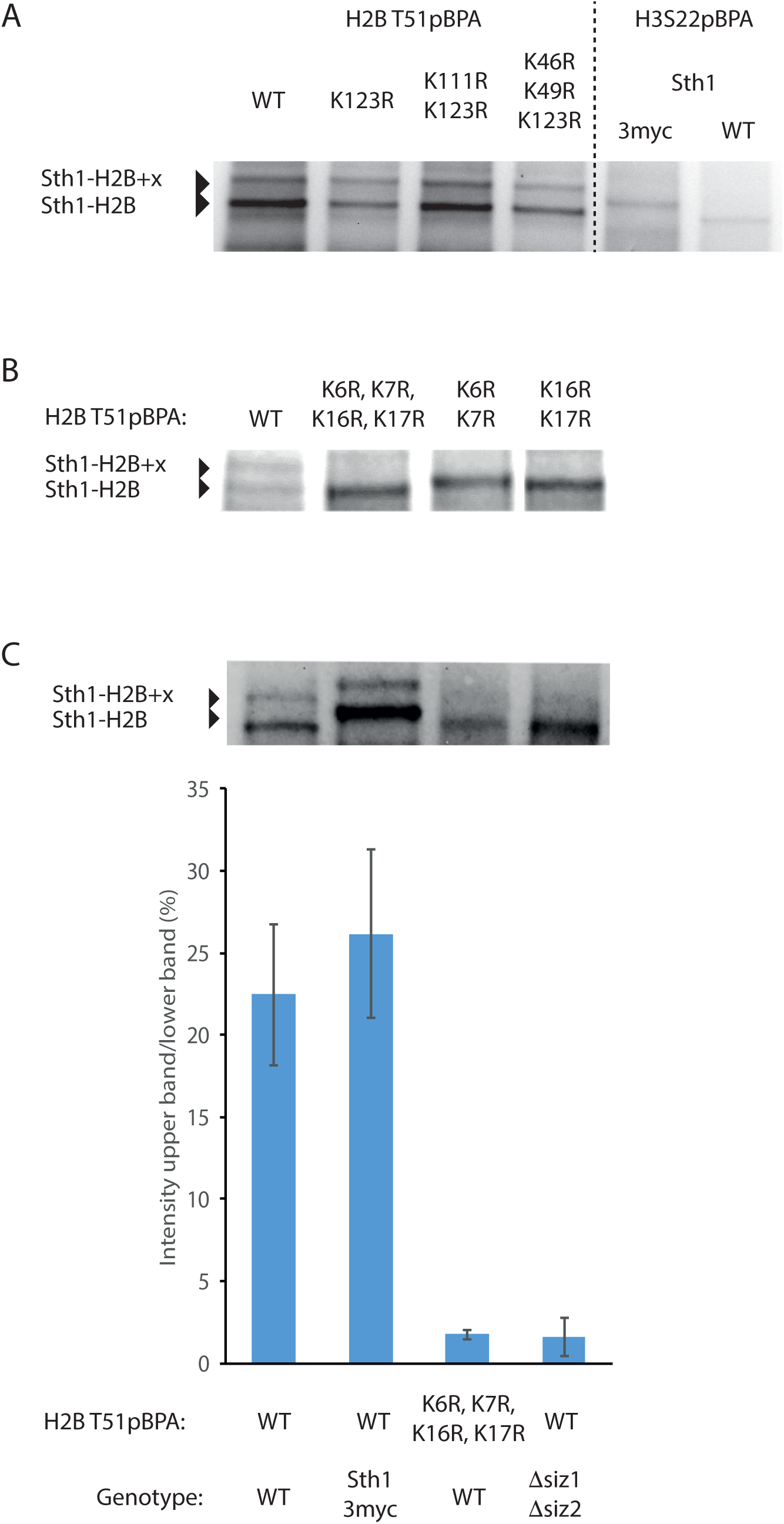
RSC prefers binding of H2B SUMOylated nucleosomes. A) Crosslink pattern of H2B T51pBPA is not affected by mutation in lysine residues reported to be subject to ubiquitinylation. Crosslinks to H3 S22pBPA are used as reference to identify Sth1-H2B crosslink. B) Effect of mutating H2B SUMOylation sites on H2B T51pBPA crosslink pattern. C) Effect of deletion of *siz1* and *siz2* on H2B T51pBPA crosslink pattern. Intensities of upper and lower crosslink bands were quantified by densitometry. Error bars are standard deviations of five independent experiments. In all panels, yeasts expressing H2B T51pBPA with the indicated mutations were UV-irradiated and whole cell lysates analysed by SDS-PAGE and Western Blot using anti-HA antibodies. Full blots in Fig. S7.

### Impact of H2B SUMOylation on RSC in vitro

In order to reveal the impact of H2B SUMOylation on the interaction of RSC with nucleosomes, we produced nucleosome core particles (NCPs) containing an in-frame fusion of SUMO to H2B (*Xenopus* sequence truncated by the first five N-terminal residues). RSC complex purified from yeast bound to SUMOylated NCPs with approximately twofold higher affinity than to unmodified NCPs in the presence of the non-hydrolysable ATP analogue AMP-PNP in biolayer interferometry (BLI) experiments (Fig. 4A&B). To further analyse RSC action on NCPs, we compared the rate of ATP hydrolysis by RSC in the presence of SUMOylated and unmodified NCPs (Fig. 4C&D). We observed a slightly increased activity on SUMOylated NCPs without a change in catalytic efficiency (V_max_/c(RSC)/K_M_). Similarly, the rate of nucleosome remodelling by RSC in electromobility shift assays was not affected by H2B SUMOylation (Fig. 4E&F). However, we observed a reduced amount of free DNA in remodelling reactions with SUMOylated NCPs, suggesting that the modification may have a modest influence on the ejection of the octamer during remodelling (Fig. 4G). Altogether our data suggest that SUMOylation of H2B *per se* only has a modest role in RSC affinity and activity *in vitro* and most likely synergizes with other factors in the recruitment of RSC to chromatin *in vivo*.

**Figure 4.**
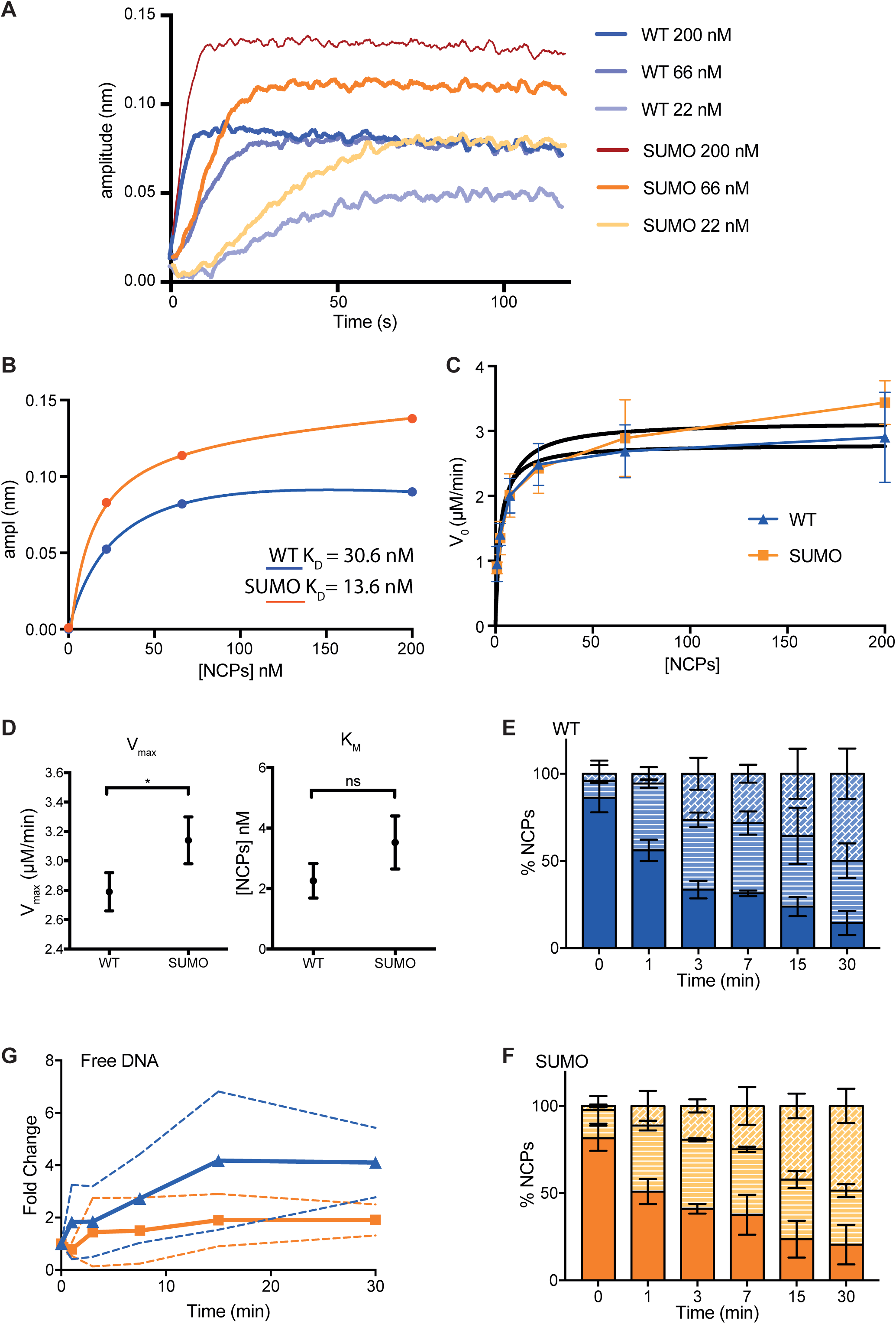
In vitro characterization of RSC-nucleosome affinity and activity. A) Biotinylated RSC complex was immobilized on streptavidin biolayer tips and NPC binding was analysed by BLI at different concentrations of NPCs (0, 22, 66, 200 nM). B) K_D_ values were determined by fitting the amplitudes of each binding kinetic with GraphPad. C) V_0_ values of RSC remodelling reactions (determined with an ADP-Glo Assay) at different concentrations of unmodified and SUMOylated NCPs (0.8, 2.5, 7.5, 22, 66, 200 nM) fit to a Michaelis-Menten equation. D) V_max_ and K_M_ determined from data shown in C for unmodified (2.79 ± 0.13 µM/min; 2.25 ± 0.56 nM) and SUMOylated NCPs (3.14 ± 0.16 µM/min; 3.52 ± 0.87 nM). E,F) RSC nucleosome remodelling activity (in the presence of Nap1) by EMSA. The relative amounts of unremodelled (top part of the bar) and two remodelled species (middle and bottom) were quantified for unmodified (WT) and SUMOylated NCPs. G) The amount of ejected DNA in EMSA was determined densitometrically and normalized to the initial amount of free DNA. Error bars are standard deviations of the means of three independent experiments. A typical EMSA gel is shown in Fig. S8.

### Modulation of Sth1-nucleosome interactions during the cell cycle

Chromatin compaction in mitosis is thought to counteract transcription by preventing access of transcription factors, RNA polymerase and chromatin remodelers to DNA [48]. To test whether RSC binding to chromatin is influenced by chromatin structure, we analysed the crosslinking efficiency of histones to Sth1 during the cell cycle (Fig. 5). Therefore, we synchronized temperature-sensitive *cdc15-2* yeasts harbouring plasmids to produce the pBPA-containing histones using a temperature shift protocol. We then sampled over time, crosslinked, and analysed the crosslink products by SDS-PAGE and Western blot. Of the four positions studied, two (H3 S22 and K56) displayed a constant crosslinking efficiency between mitosis and interphase, indicating that the RSC remodelling complex remains bound to nucleosomes throughout the cell cycle. For the other two positions (H3 T80 and H2B T51), however, we observed a reciprocal change in intensity between mitosis and interphase, indicating that chromatin structure has a subtle influence on how RSC binds nucleosomes.

**Figure 5.**
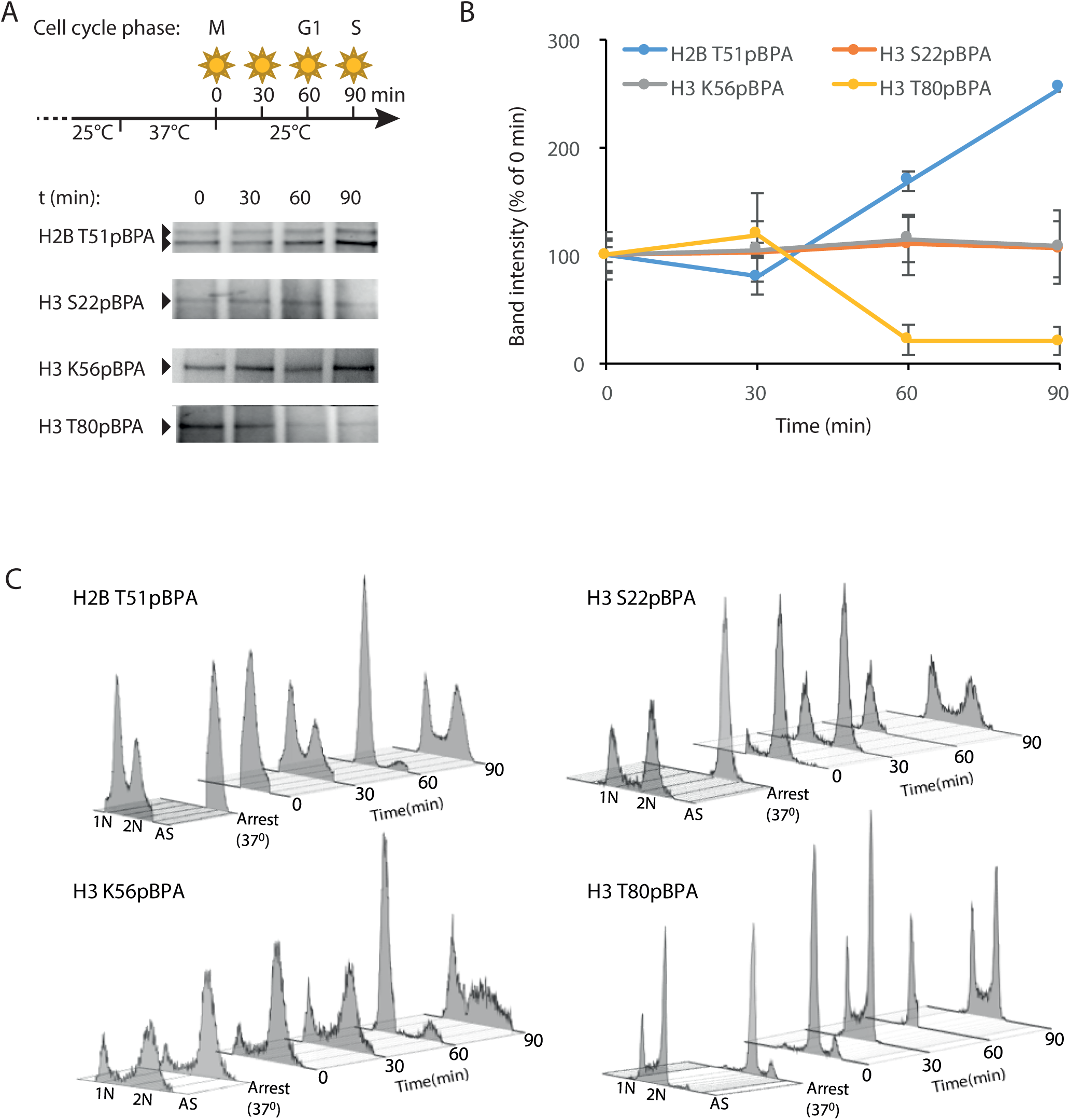
Effect of the cell cycle stage on histone-Sth1 crosslink efficiency. A) Yeasts (*cdc15-2*) expressing the indicated pBPA-containing histones were synchronized using the illustrated temperature-shift protocol. Samples were irradiated and whole cell lysates analysed by SDS-PAGE and Western blot using anti-HA antibodies. B) Band intensities of 3-4 independent replicates of the experiment shown in A were quantified by densitometry using Fuji software. Error bars are standard errors of the mean. Full blots in Fig. S9. C) FACS analysis of synchronized yeast populations analysed in A and B.

## Discussion

We analysed more than one hundred sites in core histones for their suitability for pBPA incorporation and crosslinking. Many sites gave rise to abundant and diverse crosslink products. Here, we explored whether crosslink products to Sth1 were present in the crosslinking patterns using electrophoretic mobility shift assays to reveal the footprint of the protein on the nucleosome *in vivo*. Drawing on structural information of the nucleosome-bound RSC complex [18, 19], crosslink reactions from the H3 *α*N-helix (R52, K56) most likely target the motor domain of Sth1, consistent with the observation that mutations in this helix have a strong effect on RSC remodelling activity [49].

Positions on the nucleosome surface (H2A A61, E65; H2B T51, S115; H3 T80) most likely crosslink to the SnaC domain of Sth1 [18], whereas positions in the H3-tail probably target its bromodomain in agreement with recent structural studies [22]. The RSC complex therefore contacts the acidic patches on both sides of the nucleosome simultaneously, on one side with Sth1 SnaC and with Sfh1 [18] on the other.

Our observations further show that H3 K14 acetylation is required for the interaction of the H3 tail with Sth1. Binding of the RSC complex to nucleosomes, however, is little affected by removing this mark indicating that its recruitment is controlled by additional mechanisms, e.g. general regulatory factors of transcription and DNA sequence motifs [10]. Our data suggest that the bromodomain of Sth1 binds the K14ac mark of H3 tails of neighbouring nucleosomes. This property would be well compatible with the idea that RSC contributes to the formation of a nucleosome free region at yeast promoters [10, 50].

Interestingly, position H3 T80 and H2B T51 show reciprocal changes in crosslinking intensities between mitosis and interphase (Fig. 5). We speculate that the RSC-nucleosome interaction is modulated by changes in chromatin structure during the cell cycle. Since the crosslinking efficiencies from two other sites (H3 S22 and K56) displayed hardly any changes at different cell cycle stages, we conclude that RSC activity is not controlled by eviction of the remodeler from chromatin by condensation in mitosis, as has been observed for the homologous human chromatin remodeler BRG-1 [51].

Our crosslinking experiments revealed a previously unreported preference of RSC for H2B SUMOylated nucleosomes. The crosslinking reactions from H2B positions produced a double band with the upper band being about 20-30% of the lower band intensity. In contrast, SUMOylation affects only about 5% of H2B molecules [47], implying that RSC has a strong thermodynamic preference for these nucleosomes. However, our biochemical analyses do not support this conclusion. Alternatively, RSC may be trapped kinetically at such sites by mediating the deposition of the modification through recruitment of the SUMOylation machinery. Indeed, RSC subunits were identified in a Siz1 pulldown [52]. Future experiments should address whether H2B SUMOylation modulates RSC activity *in vivo* or whether H2B SUMOylation depends on RSC activity.

## Acknowledgements

We thank Petra Geue, Simon Herrmann and Corinna Krüger for technical assistance and Andrea Musacchio for sharing reagents and his comments on the manuscript. We thank Daniela Rhodes, Fabrizio Martino and Dirk Görlich for providing plasmids.

## Extended View Figures

**Supplementary Figure 1:**
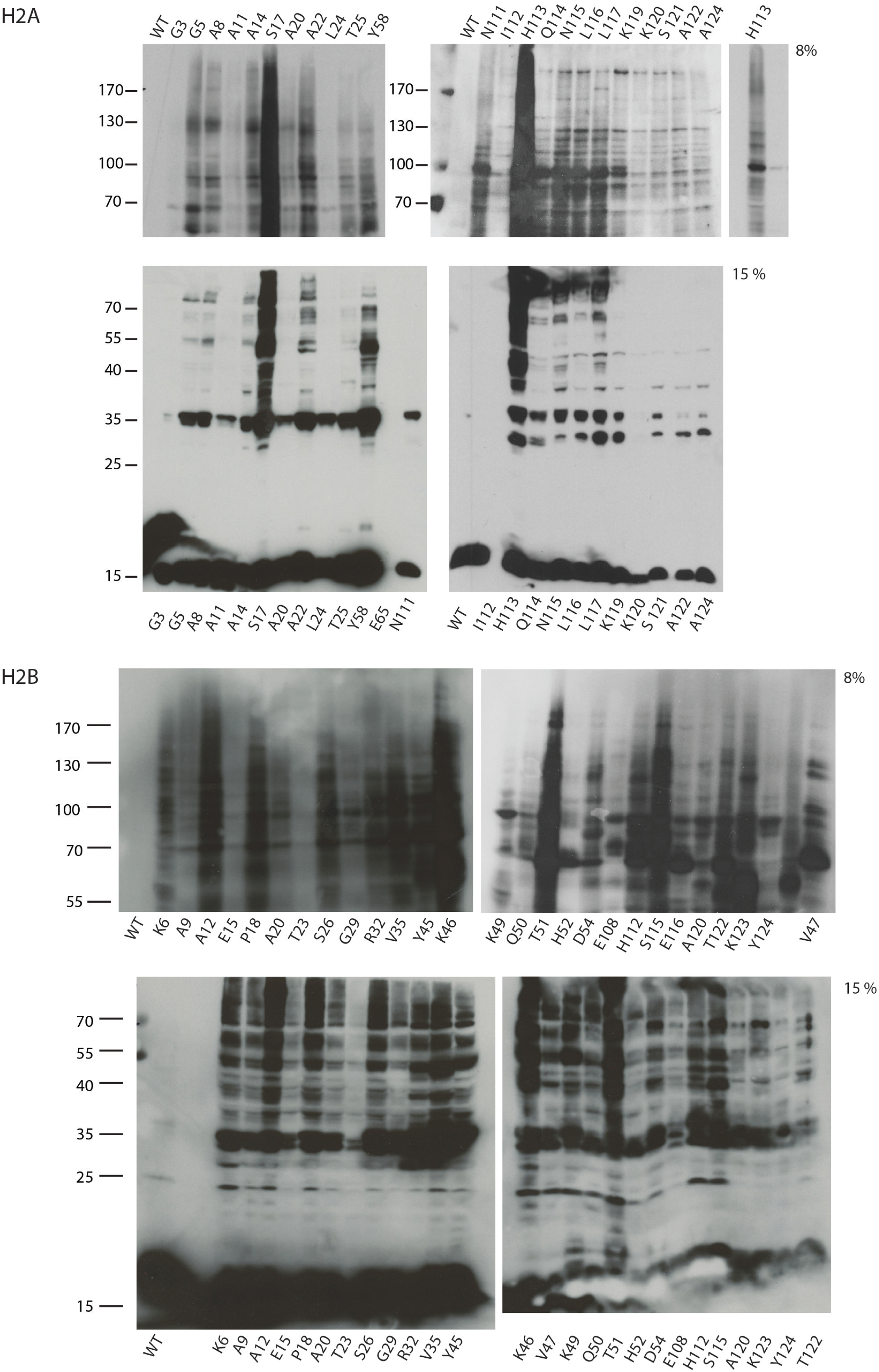
Western Blot analyses of histone H2A- and H2B-pBPA crosslink products. Yeasts expressing the indicated pBPA-containing histone were UV-irradiated and whole cell lysates analysed by SOS-PAGE and Western Blot using anti-HA antibodies.

**Supplementary Figure 2:**
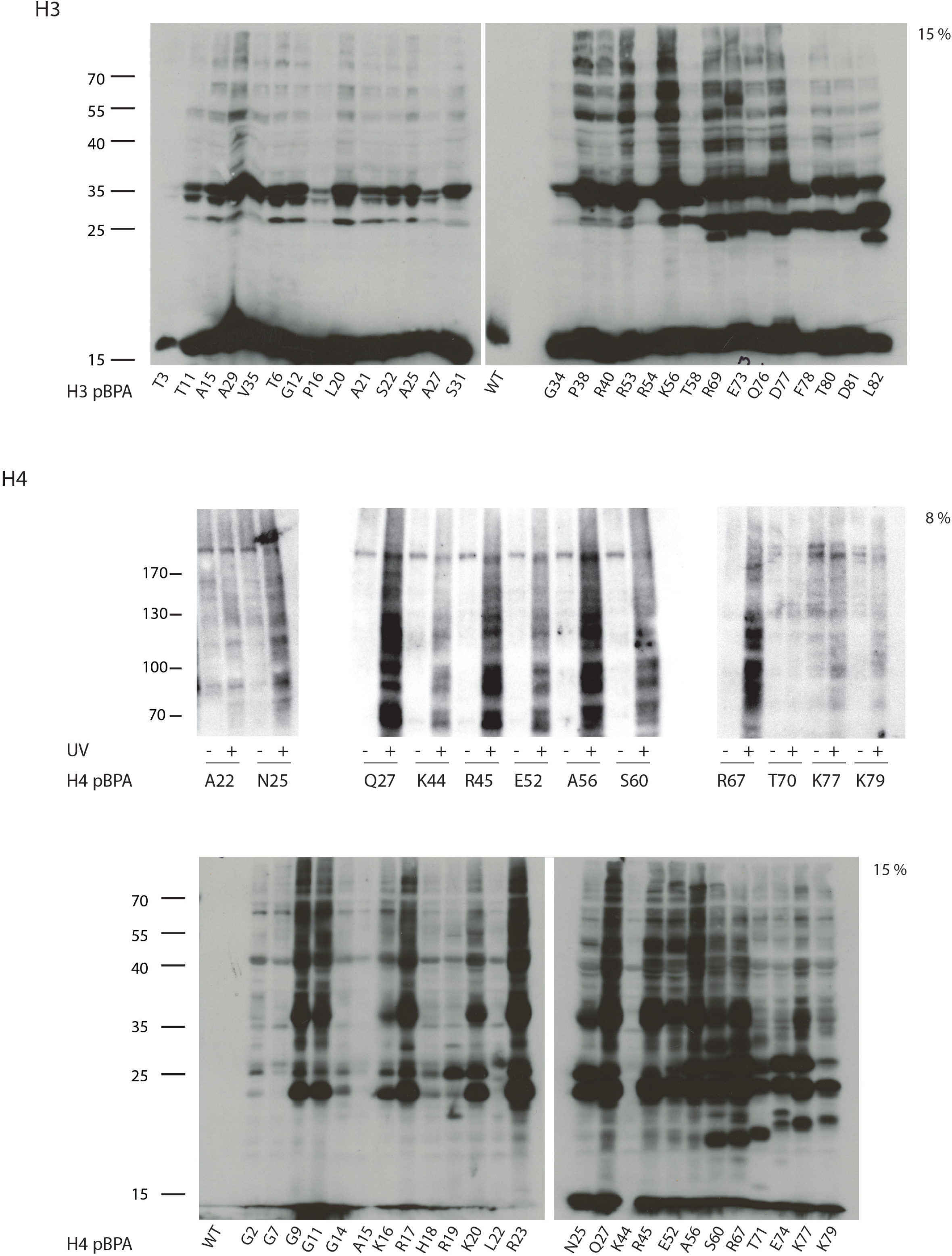
Western Blot analyses of histone H3- and H4-pBPA crosslink products. Yeasts expressing the indicated pBPA-containing histone were UV-irradiated and whole cell lysates analysed by SOS-PAGE and Western Blot using anti-HA antibodies.

**Supplementary Figure 3:**
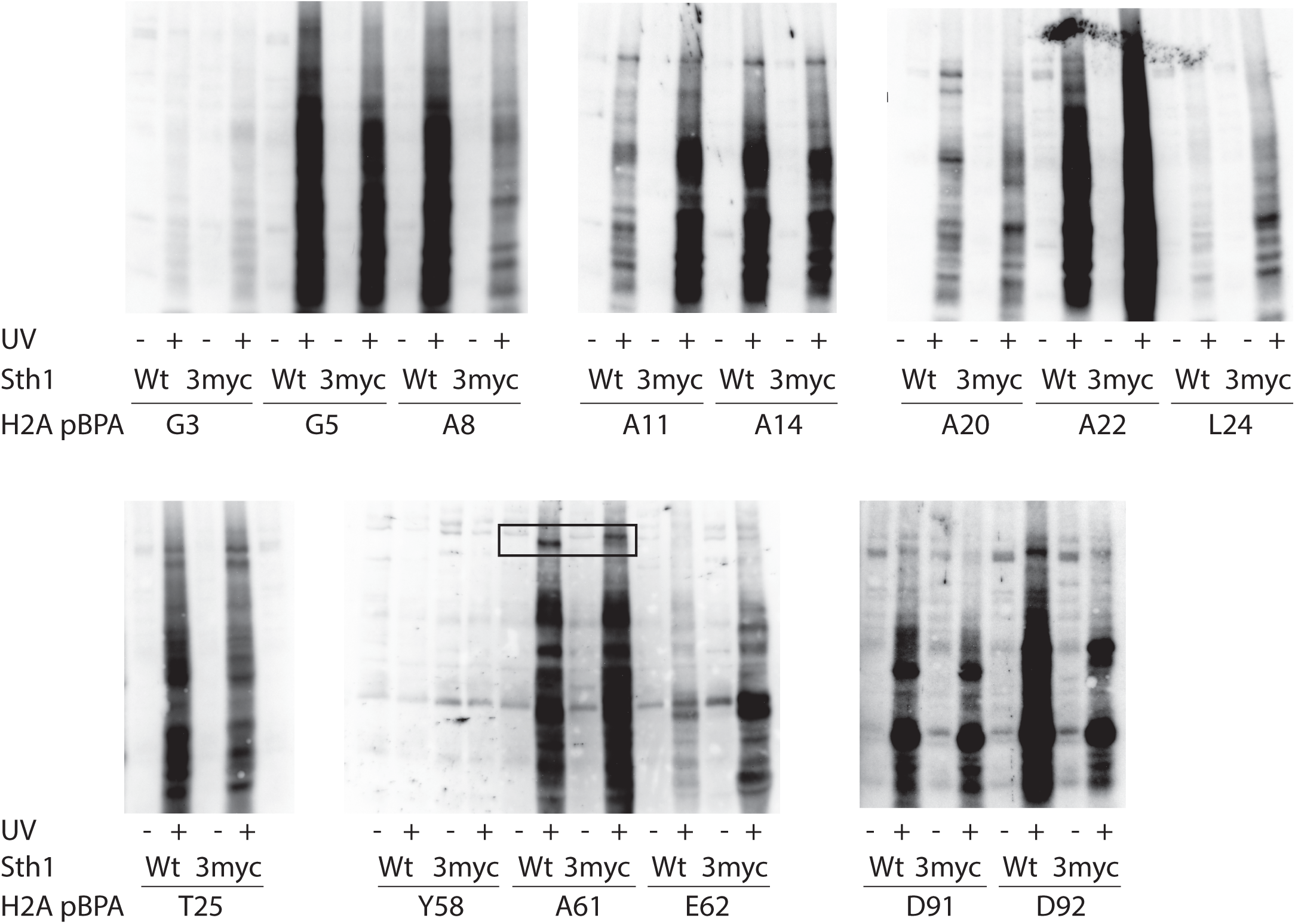
Identification of Sth1-histone H2A crosslink products. Yeasts (wild-type or Sth1-3myc) expressing the indicated pBPA-containing histone were UV-irradiated and whole cell lysates analysed by SDS-PAGE and Western Blot using anti-HA antibodies. Boxed areas show bands sensitive to Sth1-tagging.

**Supplementary Figure 4:**
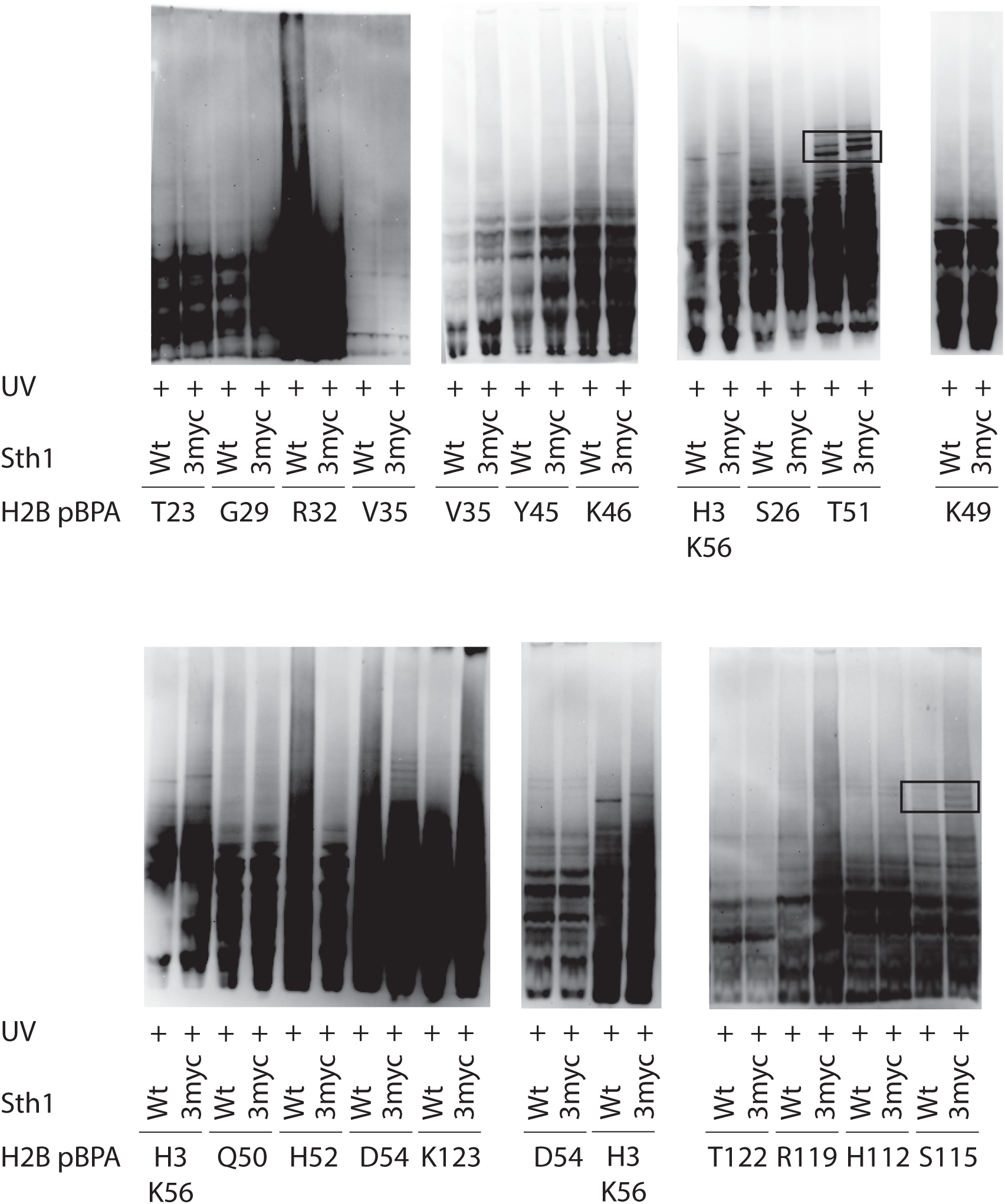
Identification of Sth1-histone H2A crosslink products. Yeasts (wild-type or Sth1-3myc) expressing the indicated pBPA-containing histone were UV-irradiated and whole cell lysates analysed by SDS-PAGE and Western Blot using anti-HA antibodies. Boxed areas show bands sensitive to Sth1-tagging.

**Supplementary Figure 5:**
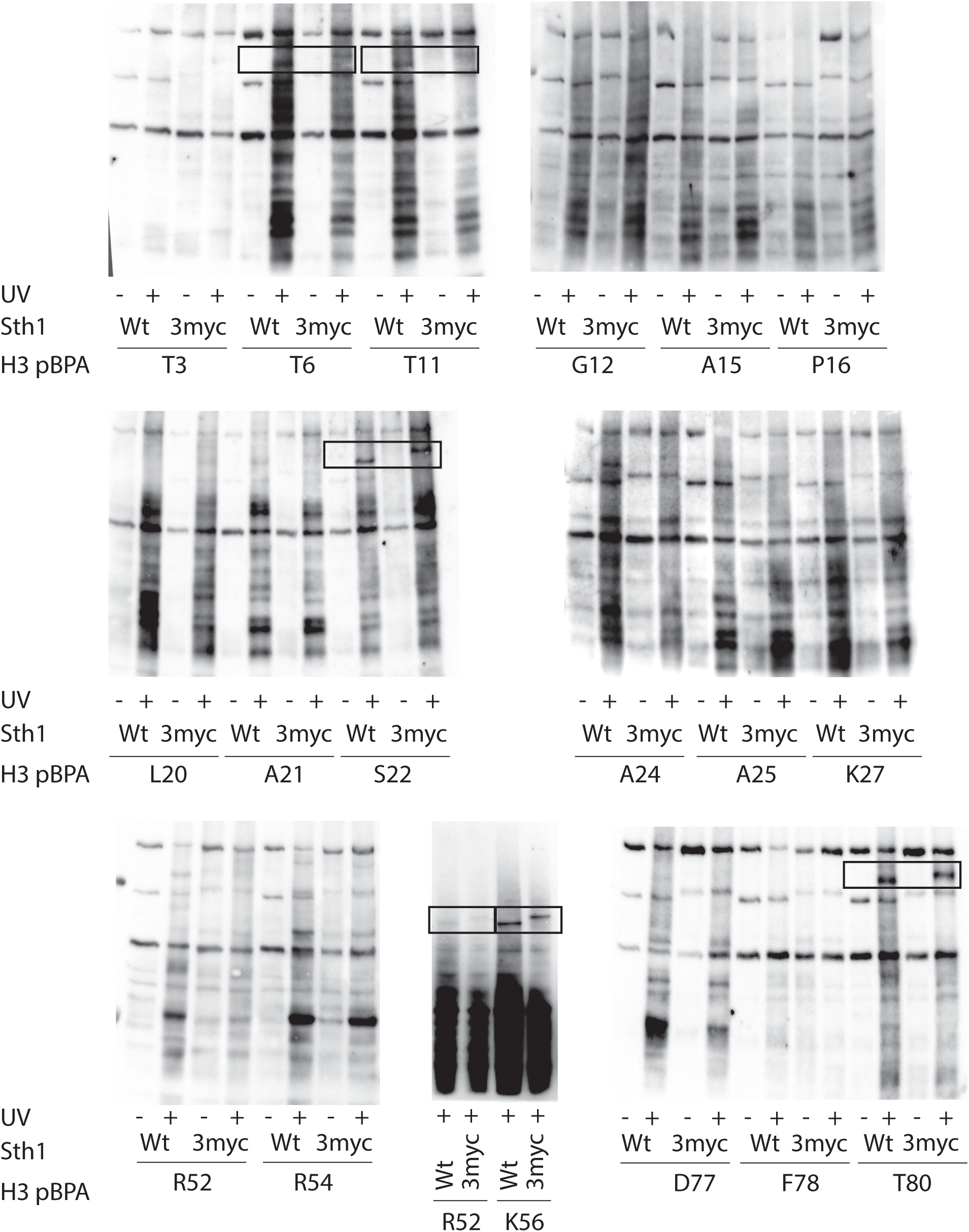
Identification of Sth1-histone H3 crosslink products. Yeasts (wild-type or Sth1-3myc) expressing the indicated pBPA-containing histone were UV-irradiated and whole cell lysates analysed by SDS-PAGE and Western Blot using anti-HA antibodies. Boxed areas show bands sensitive to Sth1-tagging.

**Supplementary Figure 6:**
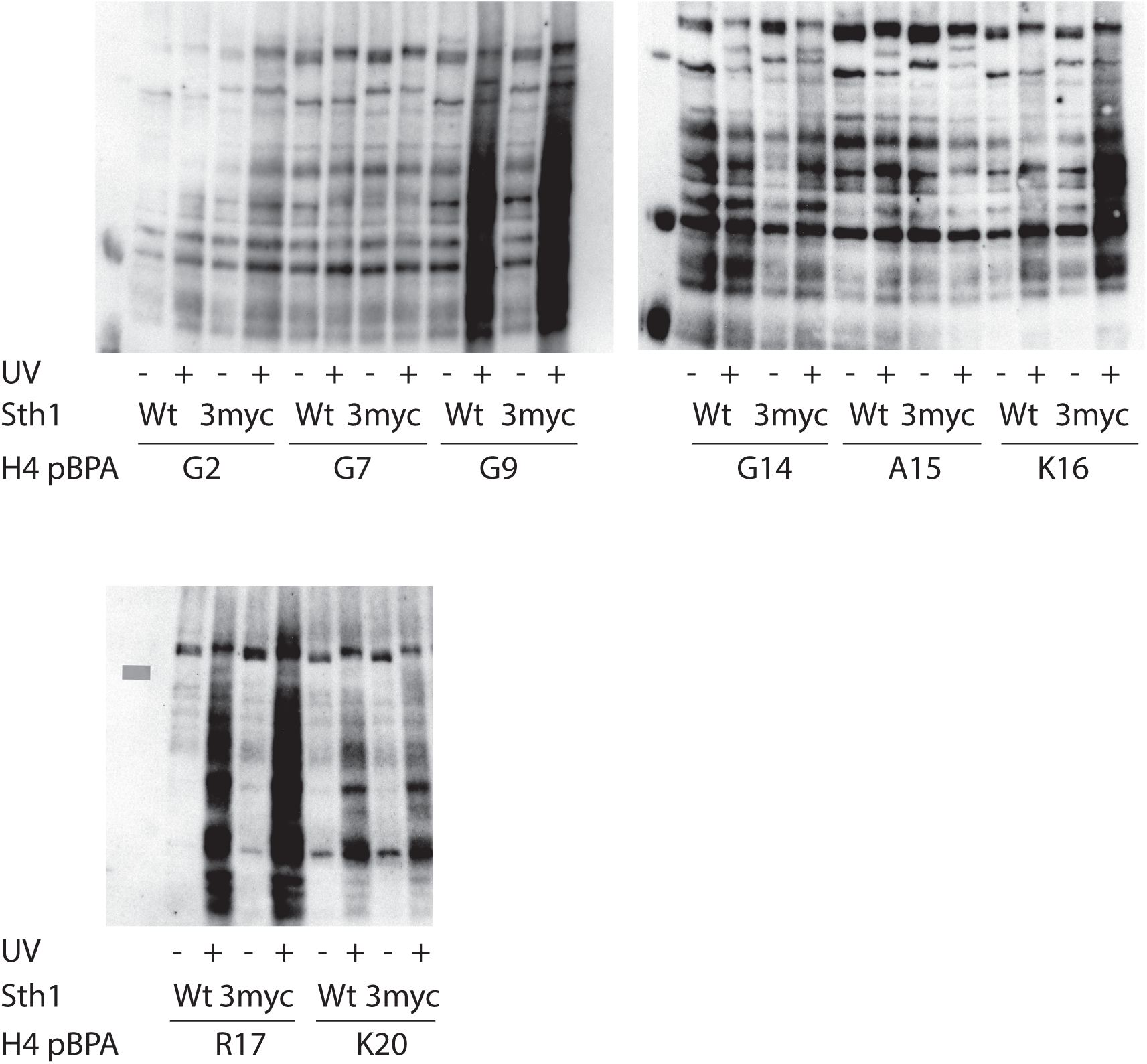
Identification of Sth1-histone H4 crosslink products. Yeasts (wild-type or Sth1-3myc) expressing the indicated pBPA-containing histone were UV-irradiated and whole cell lysates analysed by SDS-PAGE and Western Blot using anti-HA antibodies. No significant shifts were detected.

**Supplementary Figure S7:**
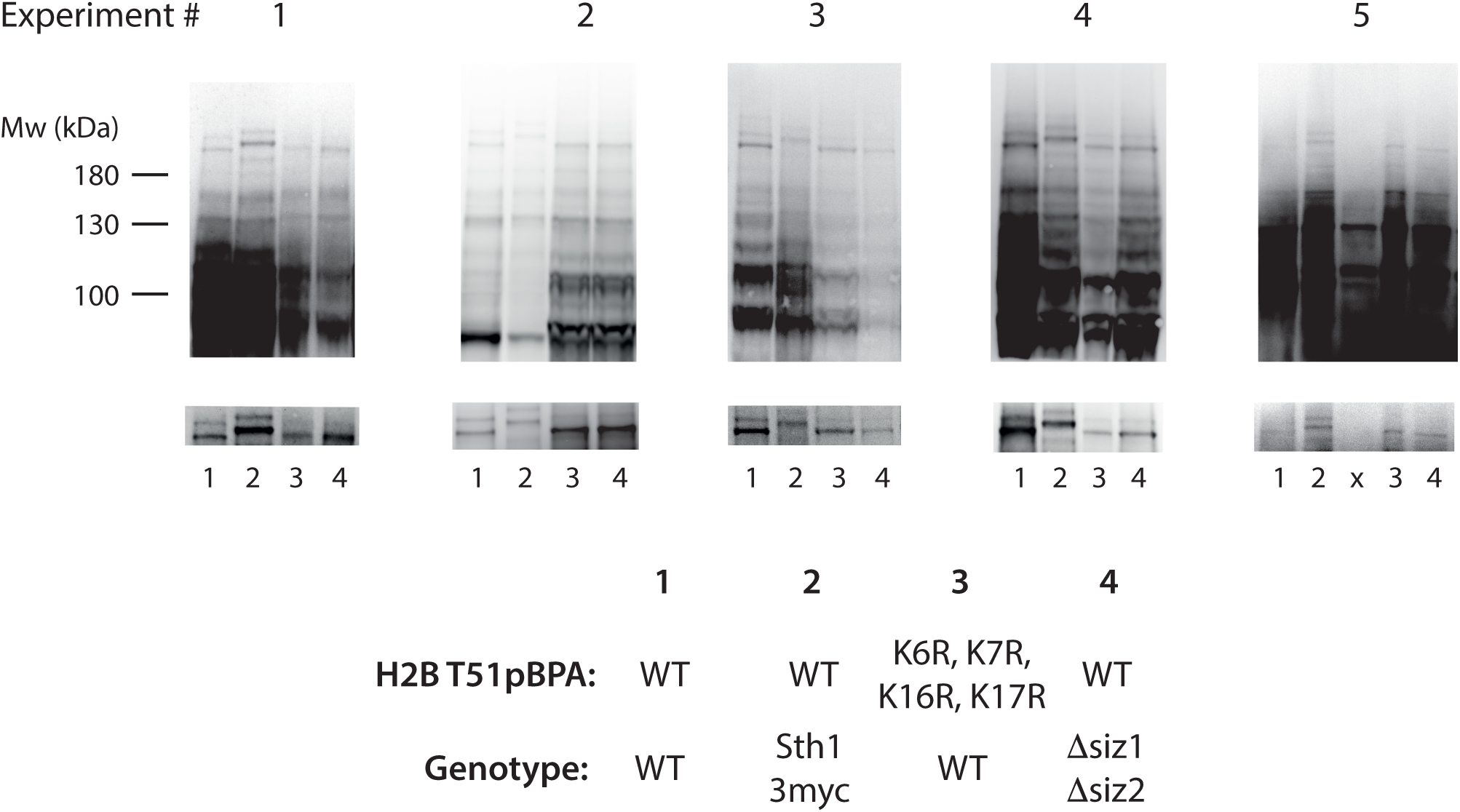
Full sized Western blots to Fig. 3

**Supplementary Figure 8:**
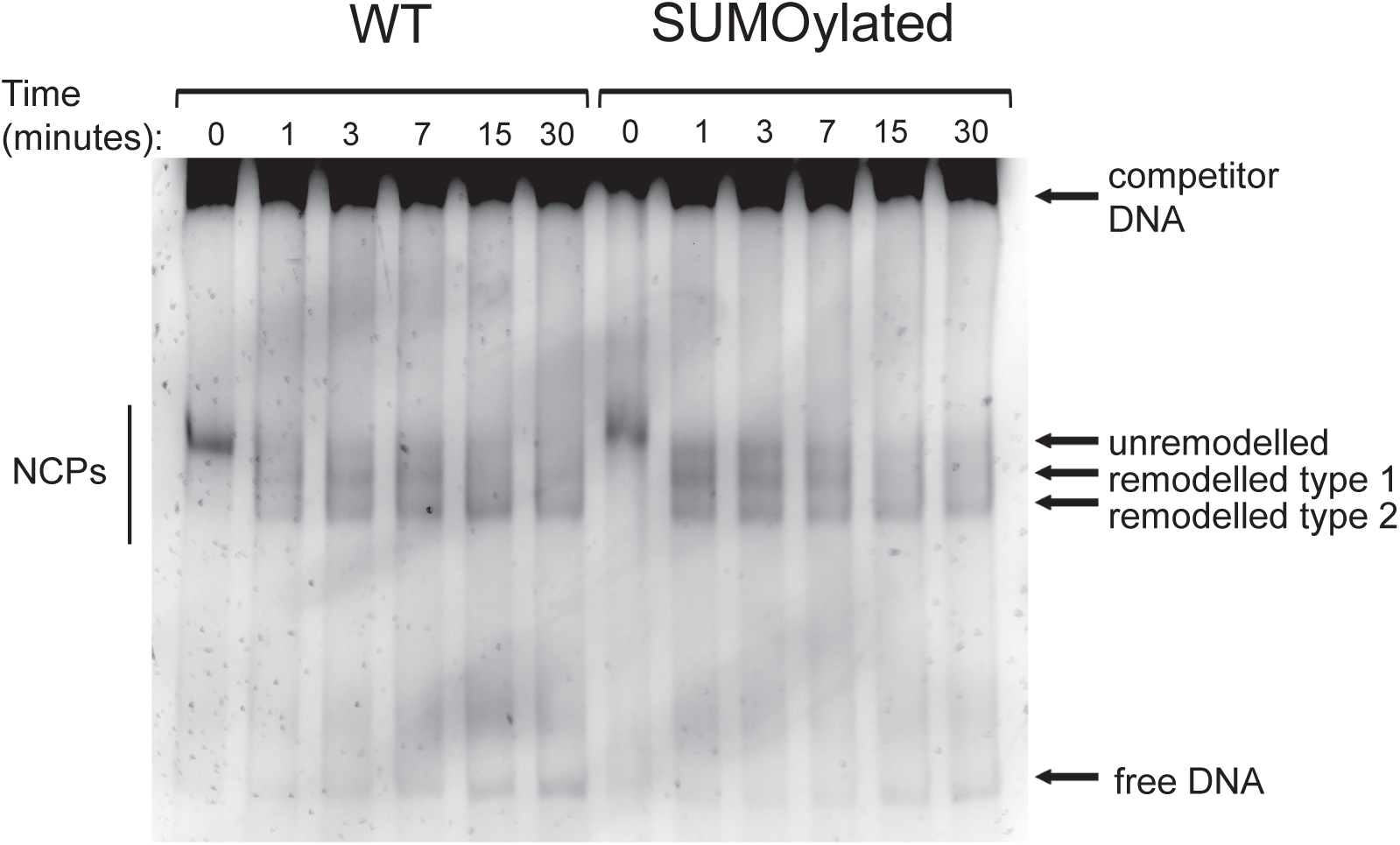
RSC remodelling reactions were analysed by EMSA. Bands for NCPs, two remodelled species and free DNA were quantified densitometrically. Data from three independent experiments are plotted in Fig. 4E-G.

**Supplementary Figure 9:**
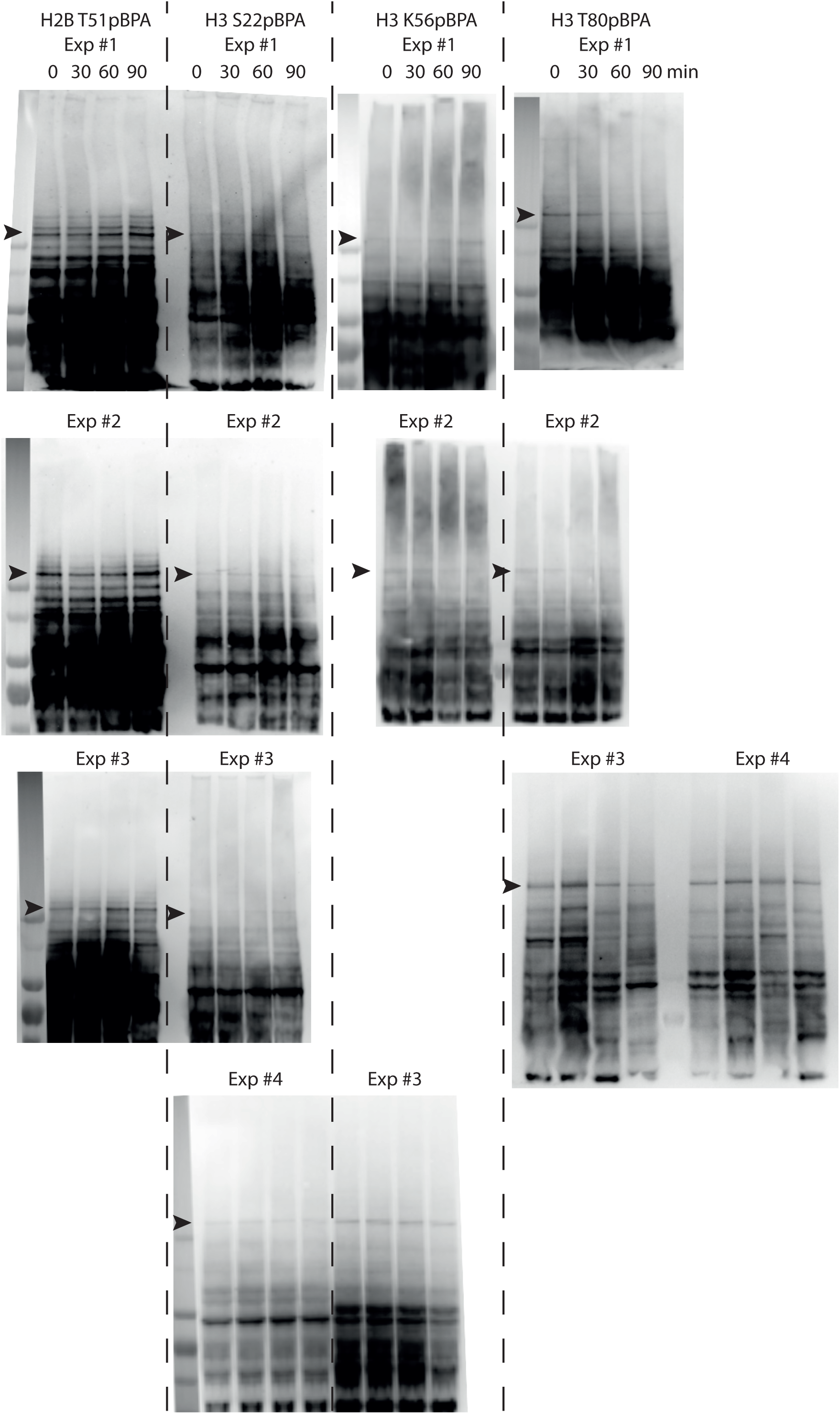
Cell cycle dependence of Sth1-histone crosslinks. Full sized Western blots to Fig. 5.

## References

1. Padeken J, Heun P (2014) Nucleolus and nuclear periphery: velcro for heterochromatin. Curr Opin Cell Biol 28: 54–60

2. Muller S, Almouzni G (2017) Chromatin dynamics during the cell cycle at centromeres. Nat Rev Genet 18: 192–208

3. Seeber A, Gasser SM (2017) Chromatin organization and dynamics in double-strand break repair. Curr Opin Genet Dev 43: 9–16

4. Talbert PB, Henikoff S (2017) Histone variants on the move: substrates for chromatin dynamics. Nat Rev Mol Cell Biol 18: 115–126

5. Tessarz P, Kouzarides T (2014) Histone core modifications regulating nucleosome structure and dynamics. Nat Rev Mol Cell Biol 15: 703–8

6. Clapier CR, Iwasa J, Cairns BR, Peterson CL (2017) Mechanisms of action and regulation of ATP-dependent chromatin-remodelling complexes. Nat Rev Mol Cell Biol 18: 407–422

7. Cairns BR, Lorch Y, Li Y, Zhang M, Lacomis L, Erdjument-Bromage H, Tempst P, Du J, Laurent B, Kornberg RD (1996) RSC, an essential, abundant chromatin-remodeling complex. Cell 87: 1249–60

8. Brahma S, Henikoff S (2018) RSC-Associated Subnucleosomes Define MNase-Sensitive Promoters in Yeast. Mol Cell

9. Floer M, Wang X, Prabhu V, Berrozpe G, Narayan S, Spagna D, Alvarez D, Kendall J, Krasnitz A, Stepansky A, et al. (2010) A RSC/nucleosome complex determines chromatin architecture and facilitates activator binding. Cell 141: 407–18

10. Krietenstein N, Wal M, Watanabe S, Park B, Peterson CL, Pugh BF, Korber P (2016) Genomic Nucleosome Organization Reconstituted with Pure Proteins. Cell 167: 709–721 e12

11. Musladin S, Krietenstein N, Korber P, Barbaric S (2014) The RSC chromatin remodeling complex has a crucial role in the complete remodeler set for yeast PHO5 promoter opening. Nucleic Acids Res 42: 4270–82

12. Spain MM, Ansari SA, Pathak R, Palumbo MJ, Morse RH, Govind CK (2014) The RSC complex localizes to coding sequences to regulate Pol II and histone occupancy. Mol Cell 56: 653–66

13. Hsu JM, Huang J, Meluh PB, Laurent BC (2003) The yeast RSC chromatin-remodeling complex is required for kinetochore function in chromosome segregation. Mol Cell Biol 23: 3202–15

14. Niimi A, Chambers AL, Downs JA, Lehmann AR (2012) A role for chromatin remodellers in replication of damaged DNA. Nucleic Acids Res 40: 7393–403

15. Rowe CE, Narlikar GJ (2010) The ATP-dependent remodeler RSC transfers histone dimers and octamers through the rapid formation of an unstable encounter intermediate. Biochemistry 49: 9882–90

16. Shim EY, Hong SJ, Oum JH, Yanez Y, Zhang Y, Lee SE (2007) RSC mobilizes nucleosomes to improve accessibility of repair machinery to the damaged chromatin. Mol Cell Biol 27: 1602–13

17. Patel AB, Moore CM, Greber BJ, Luo J, Zukin SA, Ranish J, Nogales E (2019) Architecture of the chromatin remodeler RSC and insights into its nucleosome engagement. Elife 8

18. Wagner FR, Dienemann C, Wang H, Stützer A, Tegunov D, Urlaub H, Cramer P (2019) Structure of SWI/SNF chromatin remodeller RSC bound to a nucleosome. bioRxiv: 800508

19. Ye Y, Wu H, Chen K, Clapier CR, Verma N, Zhang W, Deng H, Cairns BR, Gao N, Chen Z (2019) Structure of the RSC complex bound to the nucleosome. Science 366: 838–843

20. Zhang Y, Smith CL, Saha A, Grill SW, Mihardja S, Smith SB, Cairns BR, Peterson CL, Bustamante C (2006) DNA translocation and loop formation mechanism of chromatin remodeling by SWI/SNF and RSC. Mol Cell 24: 559–68

21. Clapier CR, Kasten MM, Parnell TJ, Viswanathan R, Szerlong H, Sirinakis G, Zhang YL, Cairns BR (2016) Regulation of DNA Translocation Efficiency within the Chromatin Remodeler RSC/Sth1 Potentiates Nucleosome Sliding and Ejection. Molecular Cell 62: 453–461

22. Chen GC, Li W, Yan FX, Wang D, Chen Y (2020) The Structural Basis for Specific Recognition of H3K14 Acetylation by Sth1 in the RSC Chromatin Remodeling Complex. Structure 28: 111-+

23. Kasten M, Szerlong H, Erdjument-Bromage H, Tempst P, Werner M, Cairns BR (2004) Tandem bromodomains in the chromatin remodeler RSC recognize acetylated histone H3 Lys14. EMBO J 23: 1348–59

24. VanDemark AP, Kasten MM, Ferris E, Heroux A, Hill CP, Cairns BR (2007) Autoregulation of the rsc4 tandem bromodomain by gcn5 acetylation. Mol Cell 27: 817–28

25. Hoffmann C, Neumann H, Neumann-Staubitz P (2018) Trapping Chromatin Interacting Proteins with Genetically Encoded, UV-Activatable Crosslinkers In Vivo. Methods Mol Biol 1728: 247–262

26. Knop M, Siegers K, Pereira G, Zachariae W, Winsor B, Nasmyth K, Schiebel E (1999) Epitope tagging of yeast genes using a PCR-based strategy: more tags and improved practical routines. Yeast 15: 963–72

27. Puig O, Caspary F, Rigaut G, Rutz B, Bouveret E, Bragado-Nilsson E, Wilm M, Seraphin B (2001) The tandem affinity purification (TAP) method: A general procedure of protein complex purification. Methods 24: 218–229

28. Guldener U, Heck S, Fielder T, Beinhauer J, Hegemann JH (1996) A new efficient gene disruption cassette for repeated use in budding yeast. Nucleic Acids Res 24: 2519–24

29. Janke C, Magiera MM, Rathfelder N, Taxis C, Reber S, Maekawa H, Moreno-Borchart A, Doenges G, Schwob E, Schiebel E, et al. (2004) A versatile toolbox for PCR-based tagging of yeast genes: new fluorescent proteins, more markers and promoter substitution cassettes. Yeast 21: 947–62

30. Spellman PT, Sherlock G, Zhang MQ, Iyer VR, Anders K, Eisen MB, Brown PO, Botstein D, Futcher B (1998) Comprehensive identification of cell cycle-regulated genes of the yeast Saccharomyces cerevisiae by microarray hybridization. Mol Biol Cell 9: 3273–97

31. Wilkins BJ, Rall NA, Ostwal Y, Kruitwagen T, Hiragami-Hamada K, Winkler M, Barral Y, Fischle W, Neumann H (2014) A cascade of histone modifications induces chromatin condensation in mitosis. Science 343: 77–80

32. Surana U, Amon A, Dowzer C, McGrew J, Byers B, Nasmyth K (1993) Destruction of the CDC28/CLB mitotic kinase is not required for the metaphase to anaphase transition in budding yeast. EMBO J 12: 1969–78

33. Haase SB (2004) Cell cycle analysis of budding yeast using SYTOX Green. Curr Protoc Cytom Chapter 7: Unit 7 23

34. Weir JR, Faesen AC, Klare K, Petrovic A, Basilico F, Fischbock J, Pentakota S, Keller J, Pesenti ME, Pan DQ, et al. (2016) Insights from biochemical reconstitution into the architecture of human kinetochores. Nature 537: 249-+

35. Shim Y, Duan MR, Chen XJ, Smerdon MJ, Min JH (2012) Polycistronic coexpression and nondenaturing purification of histone octamers. Anal Biochem 427: 190–192

36. Wittmeyer J, Saha A, Cairns B (2004) DNA translocation and nucleosome remodeling assays by the RSC chromatin remodeling complex. Method Enzymol 377: 322–343

37. Neumann-Staubitz P, Neumann H (2016) The use of unnatural amino acids to study and engineer protein function. Curr Opin Struct Biol 38: 119–128

38. Hoffmann C, Neumann H (2015) In Vivo Mapping of FACT-Histone Interactions Identifies a Role of Pob3 C-terminus in H2A-H2B Binding. ACS Chem Biol 10: 2753–63

39. Dorman G, Prestwich GD (1994) Benzophenone Photophores in Biochemistry. Biochemistry 33: 5661–5673

40. Duan MR, Smerdon MJ (2014) Histone H3 lysine 14 (H3K14) acetylation facilitates DNA repair in a positioned nucleosome by stabilizing the binding of the chromatin Remodeler RSC (Remodels Structure of Chromatin). J Biol Chem 289: 8353–63

41. Kuo MH, Brownell JE, Sobel RE, Ranalli TA, Cook RG, Edmondson DG, Roth SY, Allis CD (1996) Transcription-linked acetylation by Gcn5p of histones H3 and H4 at specific lysines. Nature 383: 269–72

42. Zhang W, Bone JR, Edmondson DG, Turner BM, Roth SY (1998) Essential and redundant functions of histone acetylation revealed by mutation of target lysines and loss of the Gcn5p acetyltransferase. EMBO J 17: 3155–67

43. Dai J, Hyland EM, Yuan DS, Huang H, Bader JS, Boeke JD (2008) Probing nucleosome function: a highly versatile library of synthetic histone H3 and H4 mutants. Cell 134: 1066–78

44. Lorch Y, Cairns BR, Zhang MC, Kornberg RD (1998) Activated RSC-nucleosome complex and persistently altered form of the nucleosome. Cell 94: 29–34

45. Robzyk K, Recht J, Osley MA (2000) Rad6-dependent ubiquitination of histone H2B in yeast. Science 287: 501–4

46. Swaney DL, Beltrao P, Starita L, Guo A, Rush J, Fields S, Krogan NJ, Villen J (2013) Global analysis of phosphorylation and ubiquitylation cross-talk in protein degradation. Nat Methods 10: 676–82

47. Nathan D, Ingvarsdottir K, Sterner DE, Bylebyl GR, Dokmanovic M, Dorsey JA, Whelan KA, Krsmanovic M, Lane WS, Meluh PB, et al. (2006) Histone sumoylation is a negative regulator in Saccharomyces cerevisiae and shows dynamic interplay with positive-acting histone modifications. Genes Dev 20: 966–76

48. Yokoyama H, Gruss OJ (2013) New mitotic regulators released from chromatin. Front Oncol 3: 308

49. Somers J, Owen-Hughes T (2009) Mutations to the histone H3 alpha N region selectively alter the outcome of ATP-dependent nucleosome-remodelling reactions. Nucleic Acids Res 37: 2504–13

50. Wippo CJ, Israel L, Watanabe S, Hochheimer A, Peterson CL, Korber P (2011) The RSC chromatin remodelling enzyme has a unique role in directing the accurate positioning of nucleosomes. Embo Journal 30: 1277–1288

51. Muchardt C, Reyes JC, Bourachot B, Leguoy E, Yaniv M (1996) The hbrm and BRG-1 proteins, components of the human SNF/SWI complex, are phosphorylated and excluded from the condensed chromosomes during mitosis. EMBO J 15: 3394–402

52. Srikumar T, Lewicki MC, Raught B (2013) A global S. cerevisiae small ubiquitin-related modifier (SUMO) system interactome. Mol Syst Biol 9: 668

